# Microglia target synaptic sites early during excitatory circuit disassembly in neurodegeneration

**DOI:** 10.1101/2024.06.13.598914

**Authors:** Alfred Yu, Li Xuan Tan, Aparna Lakkaraju, Luca Della Santina, Yvonne Ou

**Affiliations:** Department of Ophthalmology, UCSF School of Medicine, San Francisco, CA, USA; College of Optometry, University of Houston, Houston, TX, USA

**Keywords:** Microglia, Synapse, Neurodegeneration, Neuroinflammation, Excitation, PSD95, Retinal Ganglion Cell, Bipolar Cell, Synapse Disassembly

## Abstract

During development, microglia prune excess synapses to refine neuronal circuits. In neurodegeneration, the role of microglia-mediated synaptic pruning in circuit remodeling and dysfunction is important for developing therapies aimed at modulating microglial function. Here we analyzed the role of microglia in the synapse disassembly of degenerating postsynaptic neurons in the inner retina. After inducing transient intraocular pressure elevation to injure retinal ganglion cells, microglia increase in number, shift to ameboid morphology, and exhibit greater process movement. Furthermore, due to the greater number of microglia, there is increased colocalization of microglia with synaptic components throughout the inner plexiform layer and with excitatory synaptic sites along individual ganglion cell dendrites. Microglia depletion partially restores ganglion cell function, suggesting that microglia activation may be neurotoxic in early neurodegeneration. Our results demonstrate the important role of microglia in synapse disassembly in degenerating circuits, highlighting their recruitment to synaptic sites early after neuronal injury.

**Highlights:** Early after transient intraocular pressure elevation:

1. Microglia increase in number, complexity, and process movement
2. Microglia-synaptic contacts increase in the inner plexiform layer
3. Microglia-synaptic contacts increase on retinal ganglion cell dendrites
4. Microglia depletion partially restores ganglion cell function

## Introduction

High-fidelity circuit function and homeostasis are dependent on appropriate and productive interactions among diverse cell types. Microglia are the highly dynamic resident immune cells of the central nervous system (CNS) that play diverse roles in surveilling the microenvironment and respond rapidly to tissue infection and injury. In addition to their role in immunity, microglia play a role in synapse remodeling and homeostasis, especially during development, when the emergence of functional neural circuits requires the formation and elimination of synapses, a process known as synaptic pruning or elimination. Microglia-mediated synaptic pruning is critical in removing weak or supernumerary synapses during circuit refinement^1^. For example, during visual circuit development, nascent synapses are tagged with complement molecules C1q and C3, allowing microglia to recognize these excess synapses for phagocytosis^2,3^. Indeed, during development, microglia regulate laminar organization of neurons, promote cell death, limit axon outgrowth, and guide vascular networks^4^, making them crucial to circuit formation. In the disease context it has been suggested that aberrant reactivation of developmental programs of synapse pruning may result in neural circuit dysfunction^5^. Under pathological insult, microglia rapidly change their morphology and expression of signaling molecules to respond to injury. This dysregulated microglial activity can eliminate both healthy and dysfunctional synapses, impairing synaptic connectivity and ultimately circuit function^1,6^.

Synapse disassembly is an early hallmark of neurodegeneration, including during the progressive loss of ganglion cells in glaucoma, a blinding disease for which the role of retinal microglia and neuroinflammation is becoming better understood both in animal models and in human postmortem tissue^7–13^. In the retina, microglia occupy distinct regions that include the synaptic layers, the outer plexiform layer (OPL) and the inner plexiform layer (IPL). Notably, mounting evidence highlights retinal and optic nerve microglia activation as an early event in rodent models of experimental glaucoma^14–17^, although whether microglia serve a neuroprotective or neurotoxic role remains a controversial question^18–24^. Furthermore, the activation of microglia in the retinal IPL, their interaction with synapses, and their role in synapse disassembly in experimental glaucoma remain unclear.

Previously, we demonstrated that an early event in glaucomatous neurodegeneration is a reduction of synapse density on individual alpha ganglion cells prior to dendrite retraction and cell death^9^. More recently, we showed that after transient elevation of intraocular pressure (IOP), synapse loss occurs throughout the retinal IPL in a sublamina-dependent fashion^25^. Here, after transient IOP elevation, activated microglia increase in density throughout the IPL, increase their motility, and exhibit colocalization with synaptic components. Microglia increase contact with individual ganglion cell synaptic sites, including both intact synapses and disassembled synapses where the presynaptic component was absent. However, microglia depletion did not result in excitatory synapse preservation but partially restored ganglion cell function. Taken together, these experiments suggest that microglia play a role in the disassembly of synaptic connections by increasing in number after neuronal injury and colocalizing with synaptic components at excitatory synaptic sites on degenerating ganglion cell dendrites.

## Results

### After IOP elevation, microglia increase in number, complexity, and process movement in the IPL

In order to assess the role of microglia in synapse disassembly of degenerating postsynaptic neurons, we used a laser-induced ocular hypertension (LIOH) model to transiently elevate IOP in CX3CR1^GFP^ mice, which results in RGC loss but not until 14 days after IOP elevation^9,11,25,26^. Peak IOP occurs 24 h after the laser procedure, returning to baseline 5 days after treatment (Figure 1A). The IOP integral is higher in the lasered eyes (141.8 ± 8.8 mmHg □ days) vs. control eyes (115.4 ± 3.5 mmHg □ days) (p = 0.03), but there is no difference in RBPMS^+^ RGC density at this 7 day time point (Fig. 1B-C). In all experiments we examined the 7 day-post LIOH time point at which IOP has returned to baseline levels and is the earliest time point that we observe synapse loss on individual ganglion cells but no significant RGC death^9^.

**Figure 1.**
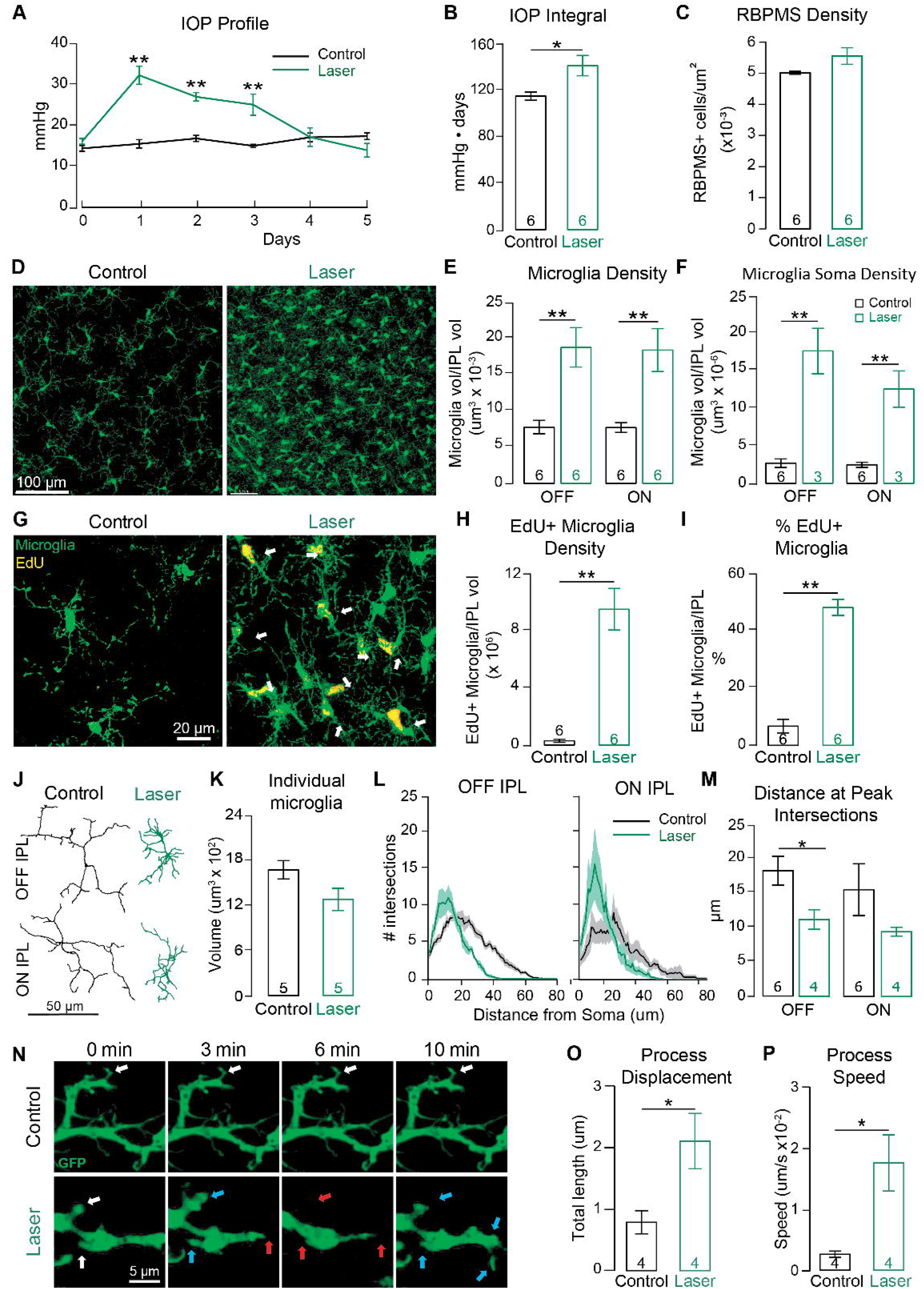
After IOP elevation, microglia increase in number, complexity, and process movement in the IPL. (A) IOP was measured daily following laser-induced ocular hypertension (LIOH), with IOP returning to baseline by day 5. (B) The lasered eye exhibited a higher IOP integral compared to the contralateral control eye. (C) Quantification of RBPMS^+^ cells shows no ganglion cell loss 7 days after IOP elevation. (D) Representative maximum z projections of microglia (green) in the IPL of control and laser conditions. (E-F) Volumetric density of microglia within the IPL is calculated in both the ON and OFF sublaminae (E) as well as (F) the density of microglia soma. (G) Representative images of EdU labeling of microglia (yellow) in control and laser conditions are quantified as (H) EdU^+^ microglia density in the IPL and (I) percentage of microglia in the IPL that are EdU^+^. (J) Representative skeletons of individual microglia. (K-M) Quantification of (K) individual microglia volume, (L) Sholl analysis of microglial processes, and (M) distance at peak intersection to determine complexity of microglia processes. Solid lines represent mean values, shaded areas represent ±SEM for control in black and laser in green. (N) Live imaging of microglia in *ex vivo* explant preparation acquired for 10 minutes; representative stills are presented here at 0, 3, 6, and 10 minutes. White arrows indicate baseline location and no change in movement. Cyan arrows indicate areas of extension and red arrows indicate areas of retraction. (O-P) Movement of microglia processes was quantified as (O) total process displacement and (P) process speed. Bars represent mean ± SEM with black and green bars representing control and laser condition respectively. The number of animals (n) is reported in each histogram. Statistics: Mann-Whitney U test; * < 0.05, ** < 0.01.

Microglia are activated in the IPL following IOP elevation as demonstrated by recruitment and proliferation of microglia, a shift in morphology, and increased motility of microglia processes. The IPL was divided into volumetric percentile bins, where 0-40% from the INL/IPL border represents the OFF sublamina and 40-100% (100% at the IPL/GCL border) represents the ON sublamina^27^. After IOP elevation, microglia volume and number are significantly elevated in the IPL in both the ON and OFF sublaminae (Fig. 1D-F, OFF sublamina LIOH 0.018±0.003 vs CTL 0.008±0.001 cells/µm^3^, p=0.008, ON sublamina LIOH 0.019±0.003 vs CTL 0.008±0.001 cells/µm^3^, p=0.005). To determine whether the increase in microglia population in the IPL is due to proliferation rather than recruitment or infiltration from the ganglion cell layer (GCL) or outer plexiform layer (OPL), an EdU (5-ethynyl-2’-deoxyuridine) cell proliferation assay was performed and ∼50% of activated microglia is the result of proliferation during the period of IOP elevation (Fig. 1G-I, LIOH 20.92±7.20 vs CTL 0.63±0.68 cells/µm^3^, p=0.005). The change in morphology of individual microglia can be appreciated in the representative skeletons (Fig. 1J), in which activated microglia are decreased in volume (Fig. 1K). Sholl analysis of individual microglia quantified the complexity of branching of microglia processes. Following IOP elevation, microglia exhibited an increase in the number of intersections as well as a decrease in the distance that the peak intersections occurred, consistent with a more amoeboid morphology in both the ON and OFF sublaminae (Fig. 1L-M). To assess microglia dynamics 7d after IOP elevation, retinas were explanted and cultured, and microglia in the ON sublamina were imaged live using spinning disk confocal microscopy. We observed rapid movement of activated microglia processes (Fig. 1N-P), with microglia process displacement (LIOH 2.14±0.46 vs CTL 0.79±0.19 µm, p=0.04) and process speed significantly increased following IOP elevation (LIOH 0.018±0.004 vs CTL 0.003±0.001 µm/s, p=0.02).

### Microglia-synaptic contacts increase in the IPL after transient IOP elevation

To determine whether microglia play a role in synapse disassembly or engulfment, we examined synaptic protein colocalization within individual microglia by immunolabeling with an excitatory postsynaptic density scaffolding protein (PSD95) and a presynaptic ribbon protein (CtBP2) (Figure 2A). We used ObjectFinder to identify all excitatory synapses colocalized inside microglia across volumes of IPL^25,28^. The population of microglia in the IPL increased engulfment of PSD95 and CtBP2 puncta, as quantified by the density of PSD95 and CtBP2 within microglia volume in the IPL (Fig. 2B-C). Dividing the IPL into the ON and OFF sublamina revealed that the increase in PSD95 and CtBP2 engulfment is occurring in both the OFF and ON sublamina of the IPL (PSD95: OFF LIOH 7.78±1.26 *10^-4^ vs CTL 2.44±0.22 *10^-4^ puncta/μm^3^, p=0.005; ON LIOH 5.73±0.10*10^-4^ vs CTL 2.71±0.23*10^-4^puncta/μm^3^, p=0.005; CtBP2: OFF LIOH 9.15±1.56 *10^-4^ vs CTL 2.82±0.024 *10^-4^ puncta/μm^3^, p=0.008; ON LIOH 9.00±1.28*10^-4^ vs CTL 4.35±0.32*10^-4^ puncta/μm^3^, p=0.005). When analyzing PSD95 and CtBP2 puncta density colocalized within each individual microglia (Fig. 2D-E), there was no significant difference between conditions (PSD95: LIOH 0.042±0.004*10^-3^ vs CTL 0.044±0.004*10^-3^ puncta/µm^3^, p=0.75; CtBP2: LIOH 0.051±0.004*10^-3^ vs CTL 0.057±0.005*10^-3^ puncta/μm^3^, p=0.18). It has been shown in hippocampus organotypic tissue culture that rather than whole synapse engulfment, microglia prune presynaptic components via trogocytosis, or “nibbling,” engulfing small (<1 micron) presynaptic components^29^. Therefore, we next examined the size of synaptic puncta colocalized inside microglia. Analysis of PSD95 puncta size showed that PSD95 engulfed within microglia have a significantly smaller average volume compared to PSD95 puncta that do not colocalize within microglia, which was true for both control and laser conditions (Fig. 2F, LIOH engulfed 0.24±0.01 vs non-engulfed 0.54±0.01 μm^3^ p=0.005, CTL engulfed 0.23±0.02 vs non-engulfed 0.52±0.01 μm^3^, p=0.005). This difference was also observed for CtBP2 inside compared to CtBP2 outside of microglia (Fig. 2G, LIOH engulfed 0.27±0.02 vs non-engulfed 0.54±0.01 μm^3^, p=0.005; CTL engulfed 0.28±0.03 vs non-engulfed 0.53±-0.01 μm^3^, p=0.005).

**Figure 2.**
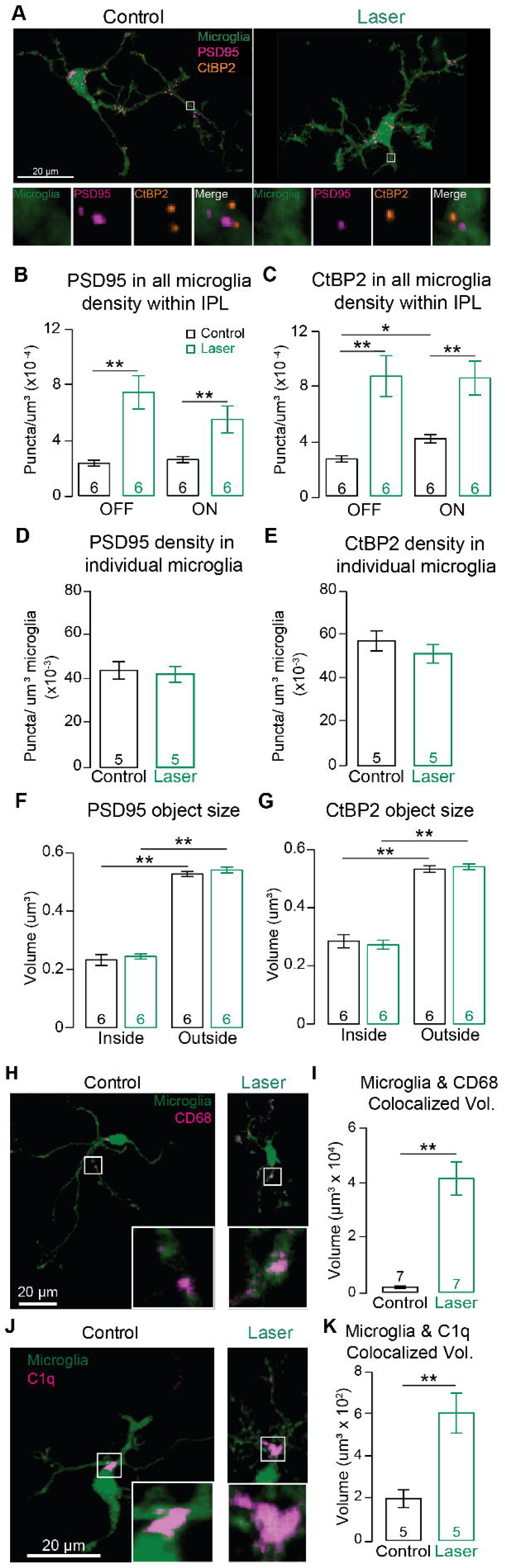
Microglia-synaptic contacts increase in the IPL after transient IOP elevation. (A) Confocal image stacks of an individual microglia (green) with engulfed PSD95 (magenta) and CtBP2 (orange) puncta in control and laser conditions. Below are zoomed in areas of the white bounding box with each signal separated and merged. (B-C) PSD95 puncta density (B) and CtBP2 density (C) in total microglia volume in the OFF and ON sublamina of the IPL. (D-E) PSD95 density (D) and CtBP2 density (E) in individual microglia volume. (F-G) PSD95 (F) and CtBP2 (G) puncta volume was analyzed inside and outside of microglia. Bars represent mean ± SEM with black and green bars representing control and laser respectively. (H) Representative images of microglia (green) and CD68 (magenta) in control and laser condition. (I) Colocalized volume between microglia and CD68. (J) Representative images of microglia (green) and C1q (magenta) in control and laser condition. (K) Colocalized volume between microglia and C1q. Zoomed in insets of the white bounding box is presented in the lower right corner of each image. Bars represent mean ± SEM with black and green bars representing control and laser respectively. The number of animals (n) is reported in each histogram. Statistics: Mann-Whitney U test; * < 0.05, ** < 0.01.

We next wanted to examine the functional phenotype of these activated microglia by assessing known markers of microglia activation and phagocytosis. CD68, a marker of microglia activation, is upregulated in microglia following IOP elevation. The colocalized volume between microglia and CD68 throughout the IPL is significantly increased (Fig. 2H-I, LIOH 41677±6130 vs CTL 1849±415 μm^3^, p=0.002). C1q, a protein complex involved in the complement pathway and known signaler of microglia for phagocytosis, is upregulated in activated microglia throughout the IPL following IOP elevation (Fig. 2J-K, LIOH: 609±95 vs CTL 200±42 µm^3^, p=0.004). Therefore, although the density of synaptic components within individual microglia was unchanged in control and LIOH conditions, activated microglia exhibited increased expression of phagocytic marker CD68 and classical complement pathway marker C1q following IOP elevation.

### Microglia target synapses on alpha ganglion cells following IOP elevation

Using the CX3CR1^GFP^ mouse along with ballistic labeling of individual RGCs and immunolabeling of synaptic components, we evaluated microglial activation and their role in synapse disassembly at the individual RGC level (Figure S1). To determine microglial interaction with RGCs, individual ganglion cells were ballistically labeled with dextran dye and microglia interaction with synaptic proteins within the ganglion cell was analyzed (Figure 3A). Alpha ganglion cells (αRGCs) were identified based on their large soma size and characteristic dendritic architecture^30^. To identify the number of contact points between microglia and individual αRGCs, we colocalized their respective signals and measured the overlapping volume, observing that both number (OFF: LIOH 41.5±4.9 vs CTL 23.2±4.1, p=0.03; ON: LIOH 37.1±7.0 vs CTL 10.1±1.8 contacts, p=0.0003) and volume (OFF: LIOH 69.4±8.2 vs CTL 26.8±4.7 μm^3^, p=0.0002; ON: LIOH 62.4±13.5 vs CTL 17.3±3.8 µm^3^, p=0.001) of contact points is increased in the laser condition (Fig. 3C-D). To determine if these contact points were at an excitatory synaptic site on the dendrite, we further colocalized the contact points to PSD95 and observed the percentage of PSD95 puncta within an individual RGC contacting microglia significantly increased in both OFF- and ON-stratifying αRGCs (Fig. 3E; OFF: LIOH 4.8±0.6 vs CTL 1.9±0.3%, p=0.0004; ON: LIOH 4.6±0.8 vs CTL 1.3±0.2%, p=0.0005). Because it has been shown that microglia prune or “trogocytose” presynaptic components in hippocampus^29^, we examined microglia colocalization with intact synapses (defined as colocalization with both pre- and postsynaptic components on the ganglion cell dendrite) and disassembled synapses (defined as colocalization with PSD95 only on the ganglion cell dendrite missing a presynaptic ribbon). When we analyzed colocalization of microglia with intact (Fig. 3F) and disassembled synapses (Fig. 3G), we observed a significant increase in colocalization with both intact and disassembled synapses in both OFF- and ON-stratifying αRGCs following IOP elevation (Intact synapses OFF: LIOH 16.8±2.2 vs CTL 6.3±0.9 puncta/μm3 (x10^-3^), p=0.0008; ON: LIOH 15.7±3.1 vs CTL 5.1±0.9 puncta/μm3 (x10^-3^), p=0.005. Disassembled synapses OFF: LIOH 5.5±0.8 vs CTL 2.5±0.5 puncta/μm3 (x10^-3^), p=0.004; ON: LIOH 5.0±1.0 vs CTL 1.3±0.2 puncta/μm3 (x10^-3^), p=0.003). In visual system development, C1q tags synapses destined for elimination^2^, thus we examined whether C1q was colocalized with both the RGC dendrite and microglia. When C1q was observed colocalized with an RGC dendrite, microglia were also present at this site, with a relationship linearly proportional to the number of C1q puncta colocalized on the RGC’s dendrite in both control and laser conditions (Fig. 3H). This data demonstrates that microglia are targeting C1q-tagged synaptic sites rather than simply making random contacts with the ganglion cell dendrites.

**Figure 3.**
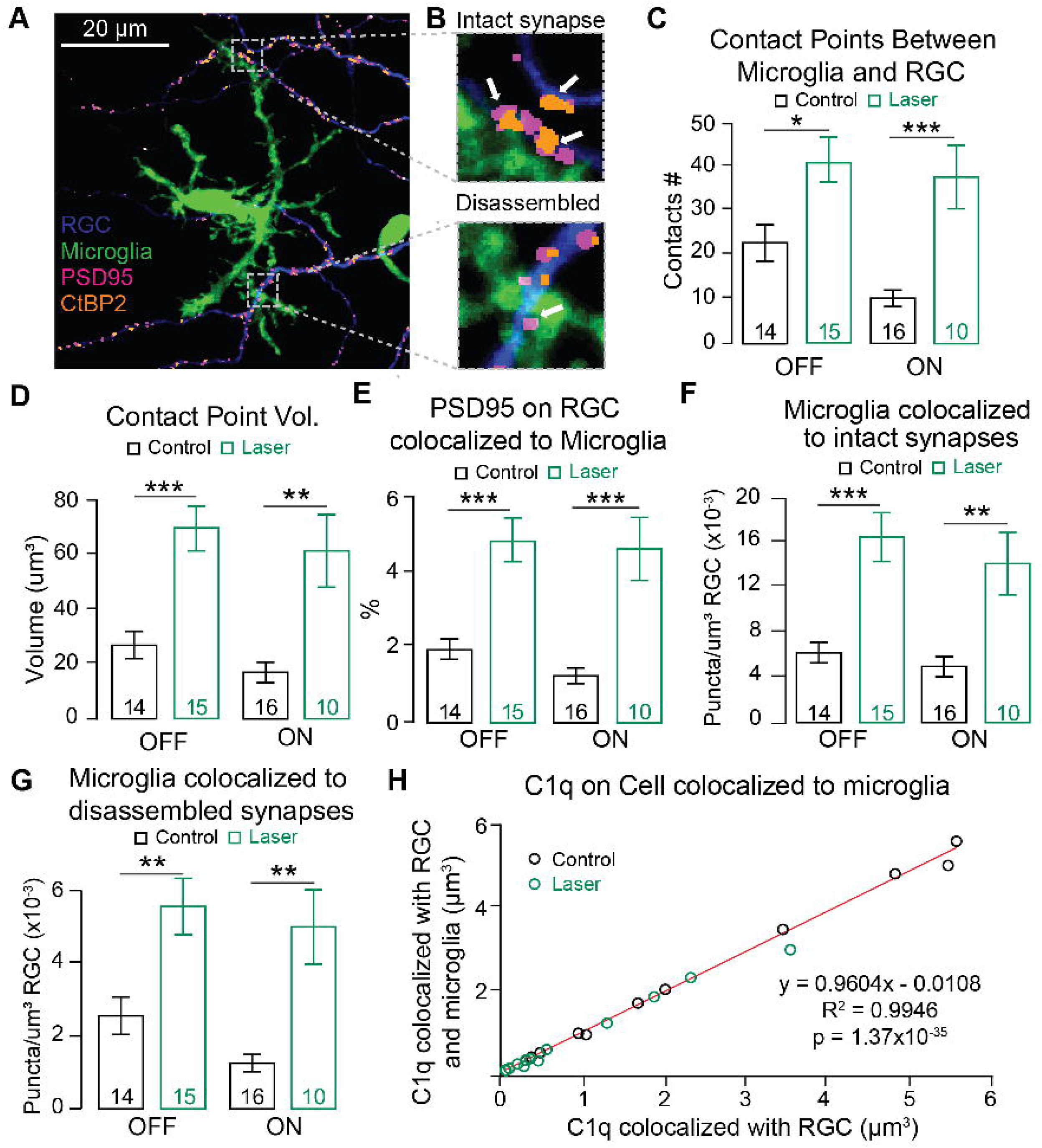
Microglia target synapses on alpha RGCs following IOP elevation. (A) Representative image of labeling of a single retinal ganglion cell (blue) with microglia (green), PSD95 (magenta), and CtBP2 (orange), see also figure S1. (B) Zoomed-in area with an intact synapse (PSD95 and CtBP2 puncta in apposition) and an area with a disassembled synapse (PSD95 with no CtBP2 puncta). (C-D) Quantification of the contact between microglia and individual cell dendrites is presented as (C) number of individual contacts and (D) the total volume of contact. (E-G) Colocalization analysis was performed to determine (E) the percentage of PSD95 on the ganglion cell colocalized with microglia; (F) the density of microglia colocalized with intact synapses on an individual ganglion cell; and (G) the density of microglia colocalized with disassembled synapses on an individual ganglion cell. (H) Correlation plot between C1q volume colocalized on RGC dendrites (abscissa) and C1q volume also colocalized with microglia (ordinata). Bars represent mean ± SEM with black and green bars representing control and laser conditions respectively. The number of cells (n) is reported in each bar from 6 mice for OFF cells and 3 mice for ON cells. Statistics: Mann-Whitney U test; * < 0.05, ** < 0.01, *** < 0.001.

### Microglia depletion partially restores RGC function following IOP elevation

To assess whether microglia play a neurotoxic or neuroprotective role on RGC structure and function after IOP elevation, we depleted microglia by administering PLX5622, a CSF1R inhibitor (Figure 4A). When PLX5622 was administered via diet we observed ∼99% depletion of microglia in the IPL in control eyes (CTL 7136±1066 vs CTL+PLX 91±99 µm^3^, p=0.01).

**Figure 4.**
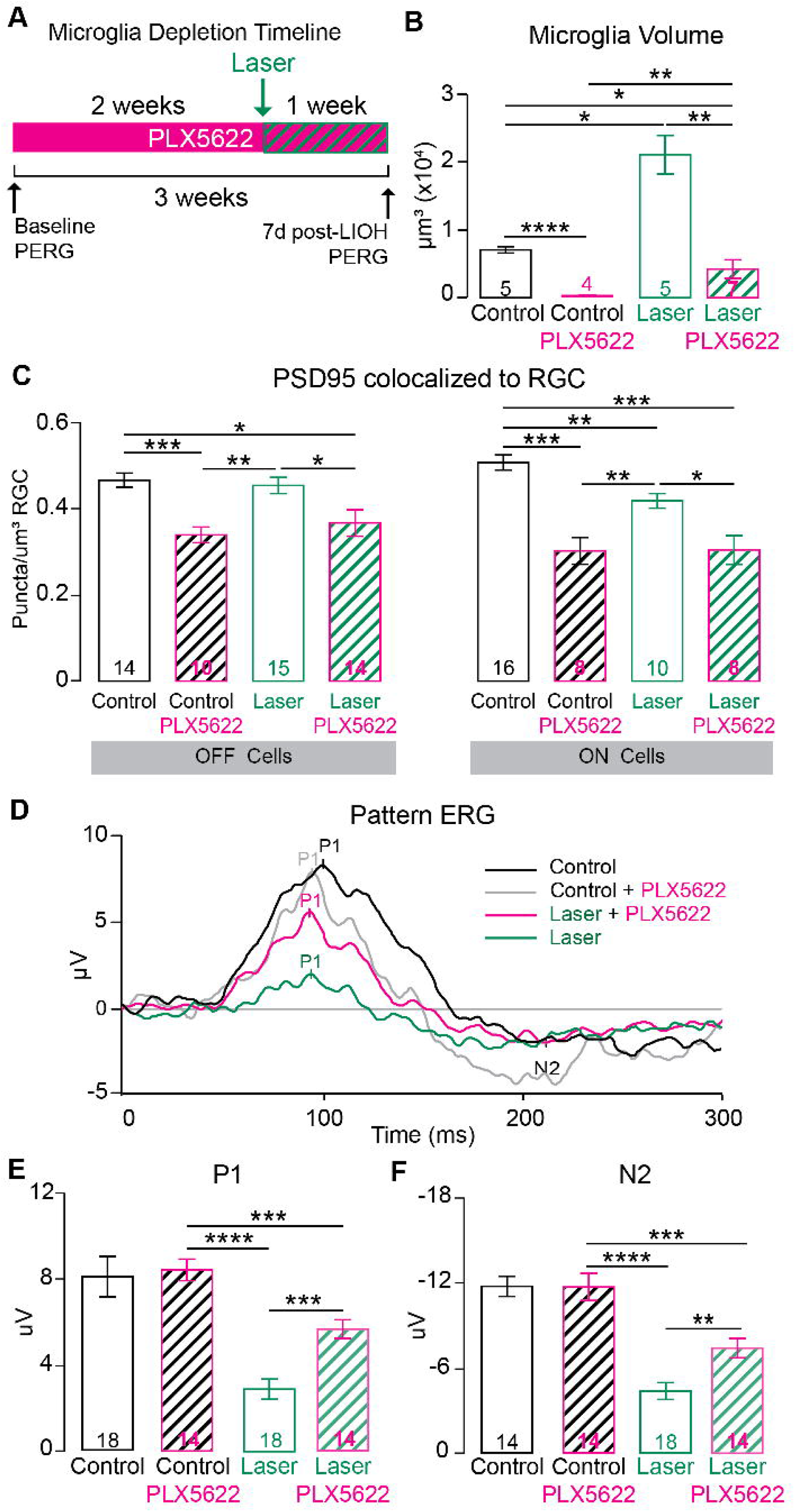
Microglia depletion partially restores RGC function following IOP elevation. (A) Experimental design of PLX5622 treatment, PERG recording, and laser-induced ocular hypertension. Following PLX5622-induced microglia depletion, (B) the volume of microglia was measured in control (black bar), control with PLX5622 treatment (magenta bar with black stripes), laser condition (green bar), and laser condition animals on PLX5622 treatment (magenta bar with green stripes). (C) Colocalization analysis of PSD95 on individual ON and OFF alpha ganglion cells is represented as synapse density across the four experimental groups. (D) Representative PERG traces of the four experimental groups. (E-F) Analysis of PERG amplitude (E) P1 and (F) N2 was quantified. Bars represent mean ± SEM. The number of animals (n) is reported in each histogram. Statistics: Mann-Whitney U test; * < 0.05, ** < 0.01, *** < 0.001, **** < 0.0001.

Lasered eyes from the same animals that were on PLX5622 diet had a significant increase in microglia volume compared to control eyes from animals on PLX5622 (CTL+PLX 91±99 vs LIOH+PLX 4258±3676 µm^3^, p=0.006), however it was decreased when compared to control eyes from animals on normal chow and significantly decreased when compared to lasered eyes from animals on normal chow (Fig. 4B, LIOH 21372±6411 vs LIOH+PLX 4258±3676 µm^3^, p=0.006). We next examined PSD95 puncta colocalized with individual ON and OFF alpha ganglion cell dendrites (Fig. 4C) and observed a significant decrease in PSD95 density in control eyes from animals on PLX5622 chow compared to control eyes from animals on normal chow (OFF PLX 0.339±0.018 vs non-PLX 0.465±0.017 puncta/µm^3^, p=0.0009; ON PLX 0.302±0.031 vs non-PLX 0.506±0.018 puncta/µm^3^, p=0.0007).

Furthermore, laser and control eyes from animals on PLX5622 diet showed comparable reductions in PSD95 density on ON and OFF αRGCs compared to laser and control eyes from animals on normal chow (Laser OFF: PLX 0.366±0.031 vs non-PLX 0.453±0.019 puncta/µm^3^, p=0.022; Laser ON: PLX 0.304±0.033 vs non-PLX 0.418±0.017 puncta/µm^3^, p=0.023). We next examined RGC function by performing pattern electroretinograms (PERGs), in which a black-white horizontal grating is presented with a square wave profile. The PERG waveform reflects inner retina activity, with a positive wave peaking at ∼80 ms (P1) and a broad wave peaking at ∼300 ms (N2)^31^. Following IOP elevation, both the P1 and N2 amplitudes of the PERG were significantly decreased (Fig 4D-F; P1 LIOH 2.92±0.47 vs CTL 8.44±0.50 µV, p=0.00001; N2 LIOH -4.51±0.61 vs CTL -12.14±0.73 µV, p=0.00001).

With microglia depleted prior to and for 7 days during and after IOP elevation there was a significant increase in P1 and N2 amplitude compared to lasered animals on normal chow (P1 LIOH+PLX 5.67±0.43 vs LIOH 2.92±0.47 µV, p=0.0005; N2 LIOH+PLX -7.66±0.71 vs LIOH -4.51±0.61 µV, p=0.003). Taken together, this data suggests that microglia may play a role in maintaining synapses in the control condition but are neurotoxic when in an activated state.

## Discussion

The role of microglia in synapse disassembly in neurodegeneration of mature circuits is not well understood. Here we show that microglia activation occurs throughout the IPL early after IOP elevation, at a time point well before ganglion cell death. Microglia increase in number, complexity, and process movement after transient IOP elevation. Activated microglia as a population in the IPL increase their colocalization with synaptic components, increase their contact with individual ganglion cells, and target synaptic sites on both ON and OFF alpha ganglion cell dendrites. Activated microglia increase expression of phagocytic and complement factor pathway proteins CD68 and C1q, but remarkably individual microglia do not exhibit an increased density of colocalized synaptic puncta. Depletion of microglia partially restored retinal ganglion cell function, despite loss of excitatory synapses, suggesting that activated microglia in the IPL are neurotoxic to RGC function early after transient IOP elevation. Taken together, the results of this study suggest that a driving factor for synapse disassembly after neuronal injury is the increased number of microglia at injury sites, which results in increased colocalization of microglia with synaptic components. Future work is needed to determine the role of microglia in active disassembly of live synapses versus engulfment of synaptic material after ganglion cell death has already occurred.

As other groups have shown in mouse models of experimental glaucoma, microglia increase in density early and throughout the retina in response to IOP elevation^14,16,32^. These studies provided a large-scale examination of changes to microglia as a result of ocular hypertension in the different layers of the retina and at the optic nerve head. Our study focuses on the IPL and individual synapses to understand the role of microglia in the mechanism of synapse disassembly between RGCs and their presynaptic partners. Synapse loss as a result of increased IOP has been shown in multiple models of experimental glaucoma^2,9,33,34^. This phenomenon is also seen in other retinal diseases; in a model of retinitis pigmentosa, microglia are activated and increase phagocytosis of postsynaptic mGluR6^35^, a process thought to be complement-mediated. Aberrant synapse loss is also an early feature of other neurodegenerative diseases such as Alzheimer’s disease, in which microglia play a key role in synapse loss^36,37^. After IOP elevation, we observed a significant increase in the number of pre- and postsynaptic puncta engulfed by microglia, driven by an increase in microglia density. The synaptic puncta colocalized inside microglia are significantly smaller in volume compared to synaptic puncta found outside of microglia, which suggests that these synaptic components colocalized with microglia have been phagocytosed or engulfed. In addition, the smaller size of engulfed puncta is consistent with microglia trogocytosis or “nibbling” of synapses, although we cannot rule out that microglia scavenge synaptic debris rather than engulfing live synapses. However, analysis of individual ganglion cells shows that the synapses colocalized with microglia are located on structurally intact dendrites. Ultimately, live imaging of individual microglia and their interactions with synaptic components is needed to define microglia function in disease conditions.

It is known that microglia in the CNS interact closely with neurons and play a role in maintenance of their function. This close contact to neurons and other glial cells allows microglia to respond quickly to stress. Here we demonstrate that upon IOP elevation, the number of microglia contacting an individual RGC significantly increases resulting in a greater number of contact points as well as increased contact volume between microglia and RGCs. Not only are microglia interacting with the individual neuron more, but this increased interaction results in greater colocalization of microglia with synaptic sites along the dendrite of the RGC, suggesting that this increased interaction is not random but targeted towards synapses destined to undergo disassembly. However, the density of synaptic components colocalized with each individual microglia does not increase after IOP elevation, suggesting that the increased synapse loss after injury is due to recruitment of additional microglia rather than simply microglia increasing their phagocytic capacity. Therefore, treatments that address the recruitment or infiltration of activated microglia, such as complement inhibition, may be an integral part of any therapeutic strategy targeting neuroinflammation.

Microglia responding to an injury signal has been shown in some contexts to be beneficial and neuroprotective; however, chronic activation of microglia is commonly associated with neurodegeneration and inhibition of microglial activation results in some neuroprotection. For example, investigators have treated animal models of glaucoma with minocycline, glucagon-like peptide-1 receptor (GLP-1R) agonists such as NLY01 or commercially available GLP-1R agonists, and complement inhibitors in order to modulate microglia function and preserve RGC survival and/or synaptic integrity^10,19,36–38^. With evidence suggesting that microglia target excitatory postsynaptic sites after IOP elevation, we tested whether depleting microglia using PLX5622 for the duration of injury would preserve RGC function. Previous work by others depleting microglia using PLX5622 has resulted in variable outcomes: in an acute model of ocular hypertension^23^ and the chronic DBA2/J mouse model^24^, microglia depletion using PLX5622 resulted in worse RGC loss, in an optic nerve crush model RGC loss was unaffected by microglia depletion^21^, and finally in an NMDA excitotoxicity model microglia depletion led to neuroprotection^22^. Here we demonstrate that when microglia are depleted before laser-induced ocular hypertension and throughout the 7 days post-laser there is a significant preservation of RGC function as assessed by PERG. However, despite the preservation of function we did not observe preservation of synapse density along individual RGC dendrites, which may be due to several limitations. We only assessed alpha RGCs, whereas PERG assesses the function of all RGCs. In the absence of microglia, other glial cells such as astrocytes may be compensating for the lack of phagocytic activity from microglia^38^. Additionally, we know that microglia play a dynamic role in neuroprotection and neuroinflammation, so depleting them may have beneficial effects in one context and time point but may be detrimental in others. Indeed, we observed that PLX5622 treatment also decreased RGC synapse density in control eyes although no decrease in PERG function was observed. Indeed, microglia have been shown to be necessary for CNS synaptic maintenance in adulthood^39^. In the retina, Wang et al. found that microglia ablation resulted in a reduction in the b-wave of the ERG and degenerative changes in outer plexiform layer synapses on electron microscopy^40^. Finally, it is also possible that in the absence of microglia, remaining synaptic connections increase their strength to compensate for decreased synapse density, resulting in partial preservation of PERG function.

After transient IOP elevation, we previously demonstrated that RGC neurodegeneration is compartmentalized, with synapse loss preceding dendrite retraction and cell death and with synapse disassembly progressing in an IPL sublamina-dependent and ordered fashion with presynaptic component loss occurring prior to or simultaneously with postsynaptic component loss^25^. However, it was not known what role microglia play in synapse disassembly. Our findings here demonstrate that activated microglia colocalize with synaptic components throughout the IPL and target synaptic sites on ganglion cell dendrites. When microglia are depleted, we observe preservation of RGC function despite no corresponding rescue in synapse density. These findings underscore the complexity of microglia function in synapse disassembly during early stages of neurodegeneration. Identifying the precise mechanisms of microglia-synapse interactions and the relationship between microglia and injured neurons is critical to the design of novel therapeutics targeting neuroinflammation in neurodegenerative diseases such as glaucoma.

## Limitations of the study

We acknowledge several limitations. First, our findings are limited to this specific experimental glaucoma model in mice, with use of the contralateral unlasered eye as control. Tribble et al. showed that microglial activation occurred in the contralateral (control) eye when ocular hypertension was induced unilaterally using magnetic microbead injection^41^.

However, analyses performed in our laboratory demonstrated that naïve eyes and contralateral eyes did not significantly differ in microglia density (data not shown). Furthermore, if microglia were appreciably activated in the contralateral eye, use of this eye as a control would not bias in favor of identifying differences between lasered vs contralateral eye. Second, the majority of data analyzed originates from confocal microscopy examination of fixed tissue and therefore the analysis is hampered by the resolution limit of this technique. Nevertheless, the reported colocalization analysis method provides an objective measurement of fluorescent signal overlap and any limitation due to the resolution of the imaging modality equally affects control and lasered eyes. In addition, ongoing work using live imaging will reveal whether microglia are scavenging synaptic debris from apoptotic cells or engulfing live synapses. Finally, microglia were depleted with PLX5622 administered via chow, which may have off-target effects as its target, the CSF1 receptor, is expressed not only by microglia but also peripheral myeloid cells^42^. PLX5622 has also been shown to decrease astrocyte gap junction coupling^43^.

## ACKNOWLEDGMENTS

The authors wish to thank Felice Dunn for helpful discussions and comments on the manuscript. We thank Yien-Ming Kuo for technical assistance. This work is supported by NIH-NEI (EY028148 to Y.O.; EY300668 to A.L.), E. Matilda Ziegler Foundation for the Blind (grant to Y.O.), BrightFocus Foundation (grants to Y.O. and A.L.), NVIDIA corporation (GPU grant to L.D.S.), Glaucoma Research Foundation (Shaffer grants to L.D.S. and Y.O.) and All May See Foundation (research grants to Y.O., L.D.S, L.X.T., and A.L.). This research was supported, in part, by the UCSF Vision Core shared resource of the NIH/NEI P30 EY002162, and by an unrestricted grant from Research to Prevent Blindness, New York, NY.

## Author Contributions

Conceptualization, Y.O. and L.D.S.; Methodology, Y.O. and L.D.S., Software, L.D.S., Formal Analysis, A.Y., Investigation, A.Y., LXT., and Y.O., Resources, A.L. and Y.O., Writing - Original Draft, A.Y. and Y.O., Writing - Reviewing & Editing, L.D.S, L.X.T., L.D.S., and Y.O., Supervision, Y.O., Project Administration, Y.O., Funding Acquisition, Y.O. and L.D.S.

## STAR Methods

### RESOURCE AVAILABILITY

#### Lead contact

Further information and requests for resources and reagents should be directed to and will be fulfilled by Yvonne Ou.

### Materials availability

The study did not generate new unique reagents.

### Data and code availability

All data reported in this paper will be shared by the lead contact upon request. All original code has been deposited at https://github.com/lucadellasantina and is publicly available as of the date of publication. DOIs are listed in the key resources table. Any additional information required to reanalyze the data reported in this paper is available from the lead contact upon request.

### Materials and Methods

#### Animals

Male and female CD-1 albino mice (Charles River; strain 022) were crossed with B6.129P2(Cg)-*Cx3cr1^tm1Litt^*/J (The Jackson Laboratory; strain 005582) to produce CD-1 albino mice with EGFP-expressing microglia. Animals were housed in animal facilities at the University of California, San Francisco. Mice were exposed to daily light cycles of 12 hour light and 12 hour dark, and given water and standard diet ad libitum except when specified. All procedures were performed as approved by the Institutional Animal care and Use Committees at the University of California, San Francisco.

#### Laser-induced ocular hypertension (LIOH)

Female and male mice 2-3 months of age were anesthetized with intraperitoneal injection of ketamine (100 mg/kg) and xylazine (10 mg/kg). Baseline IOP measurement was taken with the Tonolab rebound tonometer (Colonial Medical Co. Inc.). Each measurement was triggered with a custom foot pedal to minimize movement of the device. A total of 3 measurements per eye (each measurement is an average of 6 readings) were taken. Ocular hypertension was induced in the left eye with an endoprobe attached to a diode laser (532 nm; Lumenis) that photocoagulated the limbal and at least 3-6 episcleral vessels in the left eye (300 mW laser power, 0.5 s duration, 100 µm diameter spot size, about 80-100 total spots. The translimbal laser treatment covered 330 degrees sparing the nasal aspect and the long posterior ciliary arteries. Lubricant ophthalmic ointment was then applied to the lasered eye and the non-lasered right eye served as the control. IOP measurements were taken daily for 7 days post-laser under isoflurane at approximately the same time of day.

Gross examination of the eyes during IOP measurements were noted and eyes that exhibited corneal edema, hyphema, inflammation, and IOP > 50 mmHg were excluded from the study.

#### Ballistic labeling of individual ganglion cells

Seven days after LIOH mice were anesthetized with isoflurane and euthanized by cervical dislocation. Eyes were carefully removed and placed in oxygenated ACSF (pH 7.4) with the following components in mm: 119 NaCl, 2.5 KCl, 1.3 MgCl_2_·6H_2_O, 2.5 CaCl_2_·2H_2_O, 1 NaHPO_4_, 11 glucose, and 20 HEPES. Retinas were then dissected from the eyecup and mounted onto nitrocellulose filter paper (Millipore). Dextran-568 coated tungsten particles were prepared (Morgan & Wong, 2008) and a helium-pressured (40 psi) Helios gene gun (Bio-Rad) was used to ballistically deliver the tungsten particles onto the retinas. Within 5 minutes confirmation of cell fill was visualized on a fluorescent cell imager (Bio-Rad). The retinas were then quickly fixed with 2% paraformaldehyde for 30 min and washed twice with 1x PBS.

#### Immunohistochemistry

Fixed retinas were incubated in blocking solution (5% normal donkey serum (Jackson ImmunoResearch), 2% BSA (Sigma), and 0.05% TritonX-100 (Sigma) in 1x PBS) overnight at 4°C. Primary antibodies are then diluted in blocking solution and incubated for 4 nights at 4°C. The retinas were incubated with the following primary antibodies: mouse monoclonal anti-CtBP2 (1:500; BD Biosciences); mouse monoclonal anti-PSD95 (1:500; Neuromab); rabbit monoclonal anti-C1q (1:500; Abcam); rat monoclonal anti-CD68 (1:500; Bio-Rad).

After washing with 0.03% Triton X-100 in PBS, the appropriate secondary antibodies were incubated overnight at 4°C (Alexa, Invitrogen, 1:1000; or Dylight, Jackson ImmunoResearch, 1:1000, conjugated fluorophores). After washing again, the retinas were mounted with Vectashield (Vector Laboratories) onto glass slides.

#### EdU assay

Click-iT Plus EdU Proliferation Kit (Invitrogen) was used to determine cells that are undergoing proliferation during the 7 days post-LIOH. EdU was dosed (50 mg/mg) every day starting at day 0 to day 6 with a 5 mg/ml stock solution. After euthanasia at 7 days post-LIOH retinas were dissected and flatmounted to detect EdU+ cells.

#### Live imaging

Imaging was performed at room temperature in an Okolab humidified microenvironmental chamber on a Nikon spinning disk confocal microscope with a 60x objective (NA 1.49). The system is equipped with: Yokogawa CSU-X1 confocal spinning disk head, Nikon Eclipse Ti2-E inverted microscope, Live-SR super-resolution module, Andor iXon Ultra 888 EMCCD camera, TI2-S-SE-E motorized stage with piezo-Z for rapid Z-stack acquisition, and a laser combiner with four solid-state lasers at 405, 488, 561, and 640 nm and the corresponding band-pass emission filter sets (Chroma) loaded on a FLI high speed filter wheel. Time-lapse image stacks of microglia in the IPL were acquired every 20 seconds for a total of 10 minutes. Retinas were flattened and mounted on nitrocellulose filter paper and placed ganglion cell side face down onto a glass bottom petri dish. The petri dish is filled with oxygenated ACSF and the retina and filter paper is weighed down with a stainless steel wire to prevent drifting, with freshly oxygenated ACSF replenished after each 10 min acquisition.

#### Microglia depletion

PLX5622 (Chemgood), a CSF-1R inhibitor, was mixed in standard AIN-76A rodent diet at 1200 ppm (Research Diets) and given to the mice ad libitum. PLX5622 was administered to the mice 2 weeks prior to LIOH and they remained on the PLX5622 diet until euthanasia at 7 days post-LIOH.

#### Pattern Electroretinogram (PERG)

Inner retinal function will be assessed using noninvasive corneal PERG recording. The PERG signal is generated by presentation of a contrast reversing grating pattern, which results in a nonlinear signal that is dependent on the functional integrity of the retinal ganglion cells (RGCs) (Porciatti, 2007). Mice were anesthetized with ketamine (100 mg/kg) and xylazine (10 mg/kg) and pupils were dilated with a drop of 1% tropicamide. Genteal (Alcon) lubricating gel was used to keep the eyes from drying and used as a coupling agent during the recording. Using the Celeris ERG system (Diagnosys), a sinusoidal grating was presented to the mouse with a background light intensity of 50 cd s/m^2^ at 100% contrast and 0.155 cycles per degree. A corneal electrode is placed on the other eye as a reference. The unit is heated to 37°C to keep the mouse warm. Two sets of 300 sweeps are presented to the mouse and averaged. Once completed the PERG stimulator was placed on the other eye and the recording procedure was repeated. Positive P1 amplitude is then quantified from the N1 trough to the P1 peak; and Negative N2 amplitude is measured from the P1 peak to the N2 trough.

#### Image Acquisition and Analysis

Confocal image stacks were acquired with pixel dimensions of 0.098 x 0.098 x 0.3 µm using an upright microscope (Leica SP8) with a 63x oil immersion objective (NA 1.40). For whole IPL analysis of microglia image stacks were acquired in the central retina at the 4 leaflets of the whole mounted retina. For single cell analysis, individual alpha ganglion cells were identified and the whole IPL was imaged to determine if the dendrites stratified in the OFF or ON sublamina. To remove thermal noise from the microscope detectors, image stacks were median filtered (kernel size 1x1x1 voxels) and converted to 8-bit.

Individual alpha RGCs and all microglia were skeletonized using the filament function in Imaris (Bitplane) and binary masks were produced by fitting to their prospective signal. The IPL was then segmented from the image stack using VolumeCut. PSD95 and CtBP2 puncta were then identified using ObjectFinder and colocalization analysis with ObjectFinder identified puncta that are correlated with microglia or the individual alpha RGC.

#### Statistics

All data reported represent mean ± SEM. Statistical significance was analyzed using the Mann-Whitney U test with significance defined as p < 0.05 (Mann-Whitney *U* Test calculator (2022, February 15); retrieved from https://www.socscistatistics.com/tests/mannwhitney/default.aspx). The number of cells/animals are reported in each histogram as well as in the figure legend.

#### Key Resources Table

| Reagent or resource | Source | Identifier |
| --- | --- | --- |
| Antibodies |  |  |
| anti-CtBP2 mouse monoclonal antibody | BD Biosciences | Cat # 612044<br>RRID:AB_399431 |
| anti-PSD95 mouse monoclonal antibody | Neuromab | Cat# 73-028;<br>RRID:AB_10698024 |
| anti-CD68 rat monoclonal antibody | Bio-Rad | Cat# MCA1957 |
| anti-C1q rabbit monoclonal antibody | Abcam | Cat# ab182451 |
| DyLight 405 AffiniPure Goat Anti-Mouse IgG, Fcg subclass 1 specific | Jackson ImmunoResearch | Cat # 115-475-205;<br>RRID:AB_2338799 |
| DyLight™ 405 AffiniPure Goat Anti-Rat IgG (H+L) | Jackson ImmunoResearch | Cat# 112-475-003; RRID: AB_2338303 |
| Alexa Fluor® 647 AffiniPure Goat Anti-Mouse IgG, Fcy subclass 2a specific | Jackson ImmunoResearch | Cat# 115-605-206; RRID: AB_2338917 |
| Other reagents |  |  |
| Click-iT™ Plus EdU Cell Proliferation Kit for Imaging, Alexa Fluor™ 555 dye | Invitrogen | Cat# C10337 |
| Dextran, Alexa Fluor™ 568; 10,000 MW, Anionic, Fixable | Invitrogen | Cat# D22912 |
| PLX5622 | Chemgood | Cat# C-1521 |
| Experimental models |  |  |
| B6.129P2(Cg)-Cx3cr1tm1Litt/J | The Jackson Laboratory | Strain #:005582;<br>RRID:IMSR_JAX:005582 |
| CD-1 albino mice (CrI:CD1(ICR) Outbred) | Charles River Laboratories | RRID:IMSR_CRL:022 |
| Software and algorithms |  |  |
| Imaris | Bitplane | RRID:SCR_007370 |
| VolumeCut | https://github.com/lucadellasantina/VolumeCut | https://doi.org/105281/zenodo.5048331 |
| ImageJ | NIH | RRID:SCR_003070 |
| ObjectFinder | https://github.com/lucadellasantina/ObjectFinder | https://doi.org/105281/zenodo.4767847 |
| Mann-Whitney U Calculator | Social Science Statistics | https://www.socscistatistics.com/tests/mannwhitney/defa |
|  |  | ult.aspx |

---

**Supplemental Figure S1.**
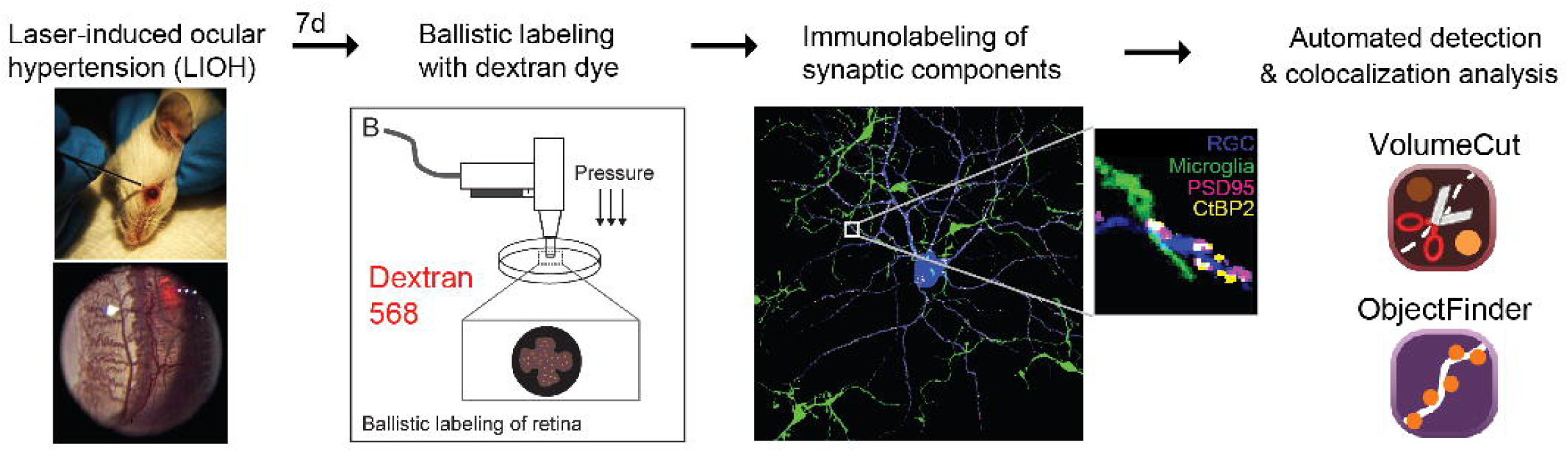
Pipeline for labeling and analysis of colocalization between microglia and individual RGCs. Laser-induced ocular hypertension (LIOH) is induced unilaterally by episcleral and limbal vessel cauterization. 7d after LIOH, individual ganglion cells are ballistically labeled with dextran dye. After immunolabeling with synaptic components, VolumeCut and ObjectFinder are used to perform semi-automated detection and colocalization analysis of synaptic components with microglia and individual alpha ganglion cells.

